## Supplemental Materials for "High-resolution neural recordings improve the accuracy of speech decoding"

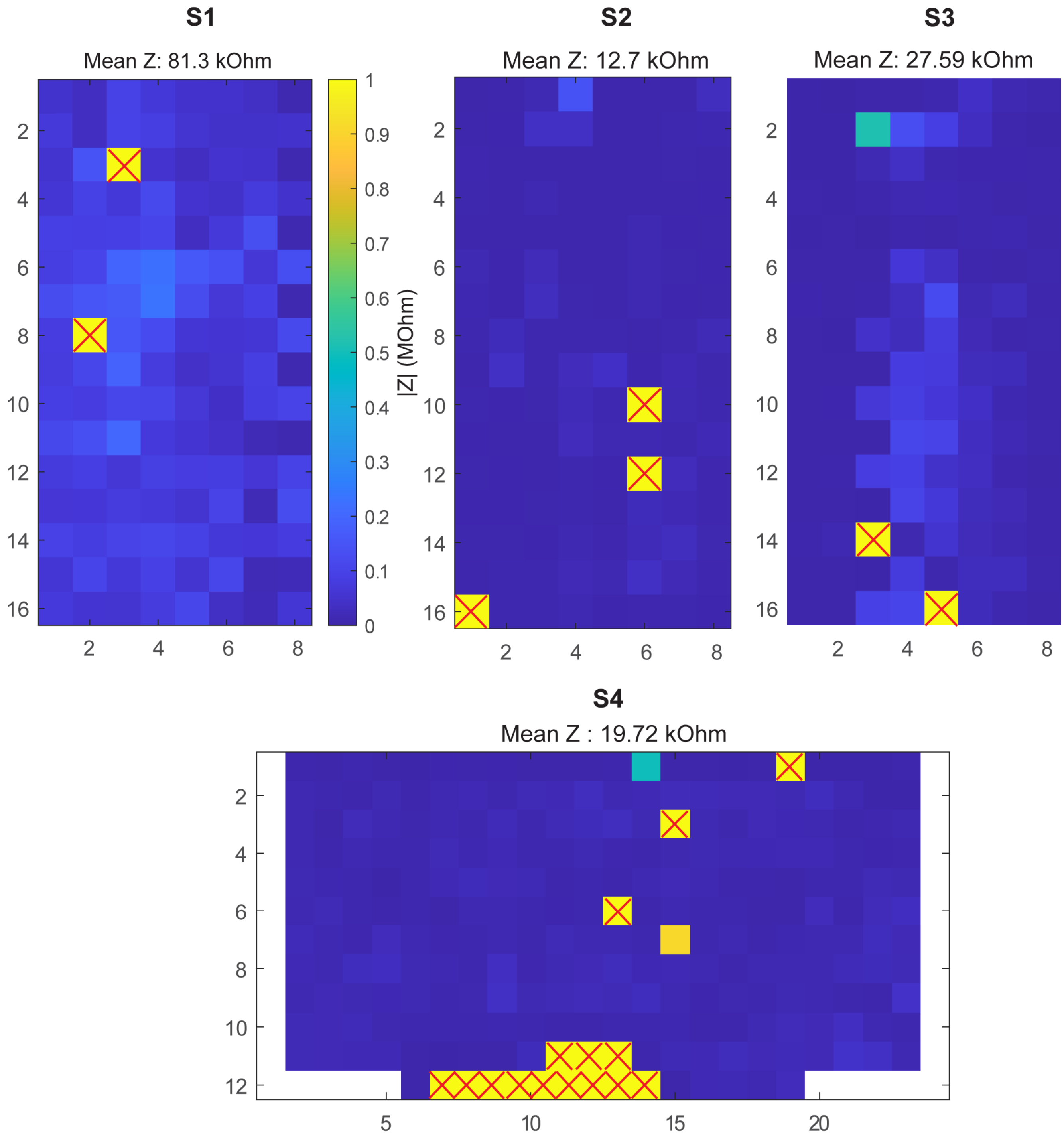


**Supplementary Figure 1**. **Spatial maps of *in vivo* electrode impedance during intraoperative µECoG implantation**. Uniform impedance measurement at 1kHz for 128-channel (S1, S2, and S3) and 256-channel (S4) LCP-TF electrode arrays demonstrate our successful implantation procedure in the intraoperative setting. Channels with impedance >1 MΩ were considered non-functional and are identified with red markers in the spatial maps.

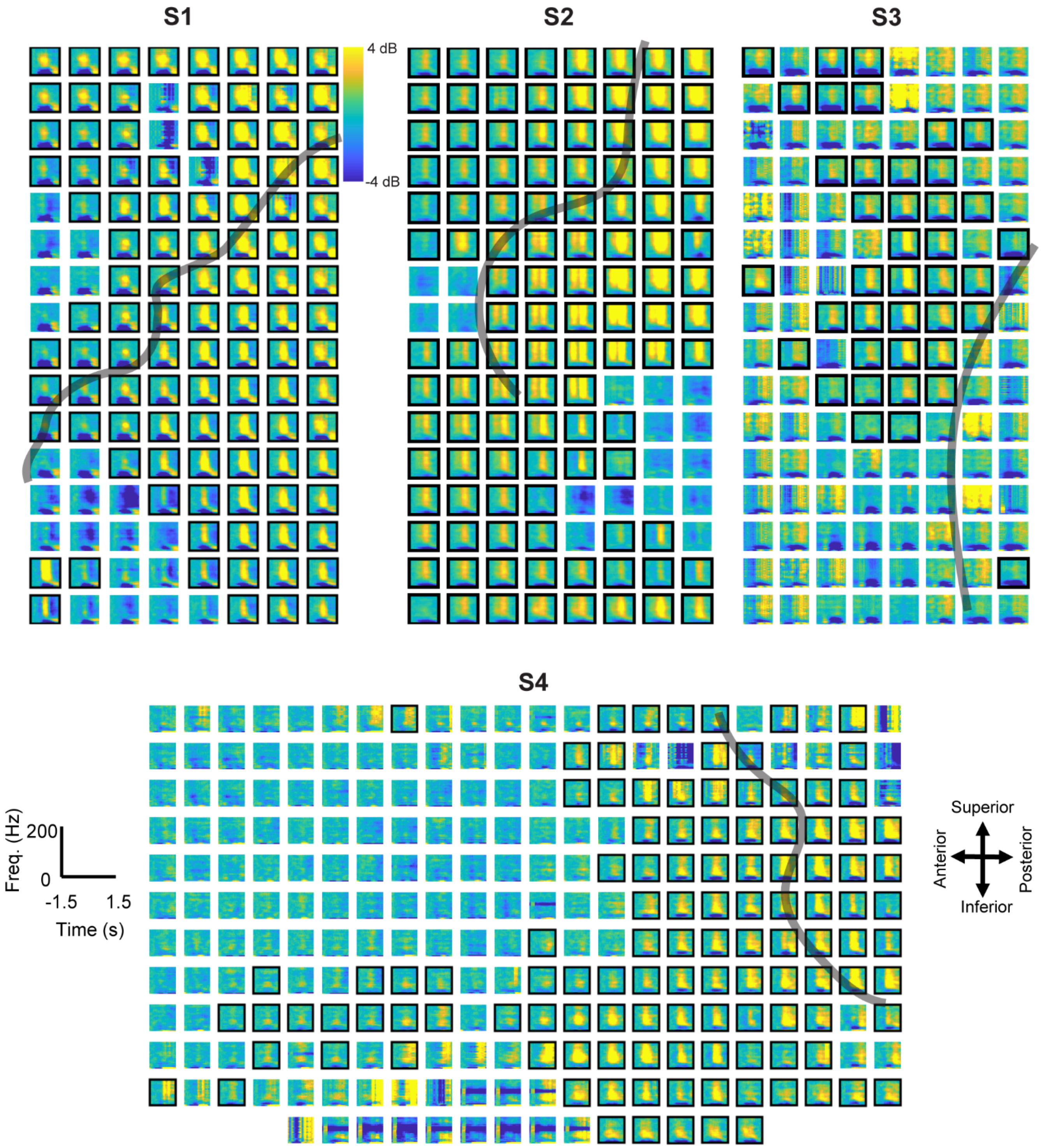


**Supplementary Figure 2**. **Spectrogram activations for four intraoperative subjects during a speech production task**. Normalized spectrograms reveal distinct spectro-temporal modulation for each μECoG channel. We focus on the significant rise in HG power during speech production (black border: one-sided permutation test, $p < 0.01$; corrected via false discovery rate method). Shaded gray curves indicate rough estimate of central sulcus from average MNE brain. Channels with significant HG modulation reveal the spatial activation profile across the μECoG array for all subjects (S1: 111/128, S2: 111/128, S3: 60/128, and S4: 149/256).

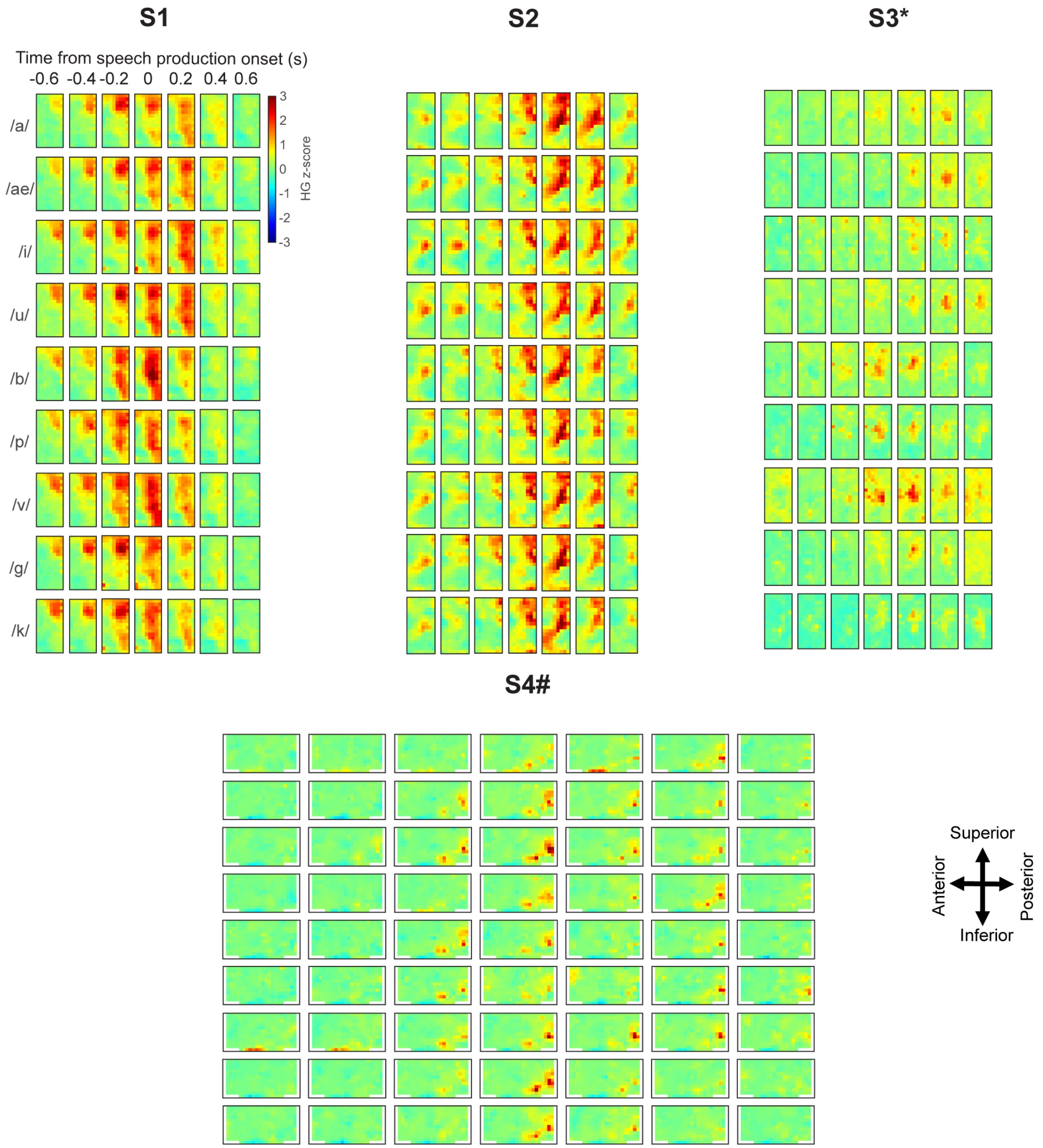


**Supplementary Figure 3**. **Fine-scale spatiotemporal activations of HG power during speech production tasks**. Channel map activations reveal distinct, highly-resolved spatio-temporal activity for all subjects during each phoneme utterance (first-position phonemes). Phonemes with similar articulatory features exhibit similar activation patterns at speech utterance. S1 and S2 exhibited stronger HG-SNR revealing clear phoneme-specific activation patterns. S3* and S4# exhibited weaker yet significant activations due to low SNR and recording duration, respectively.

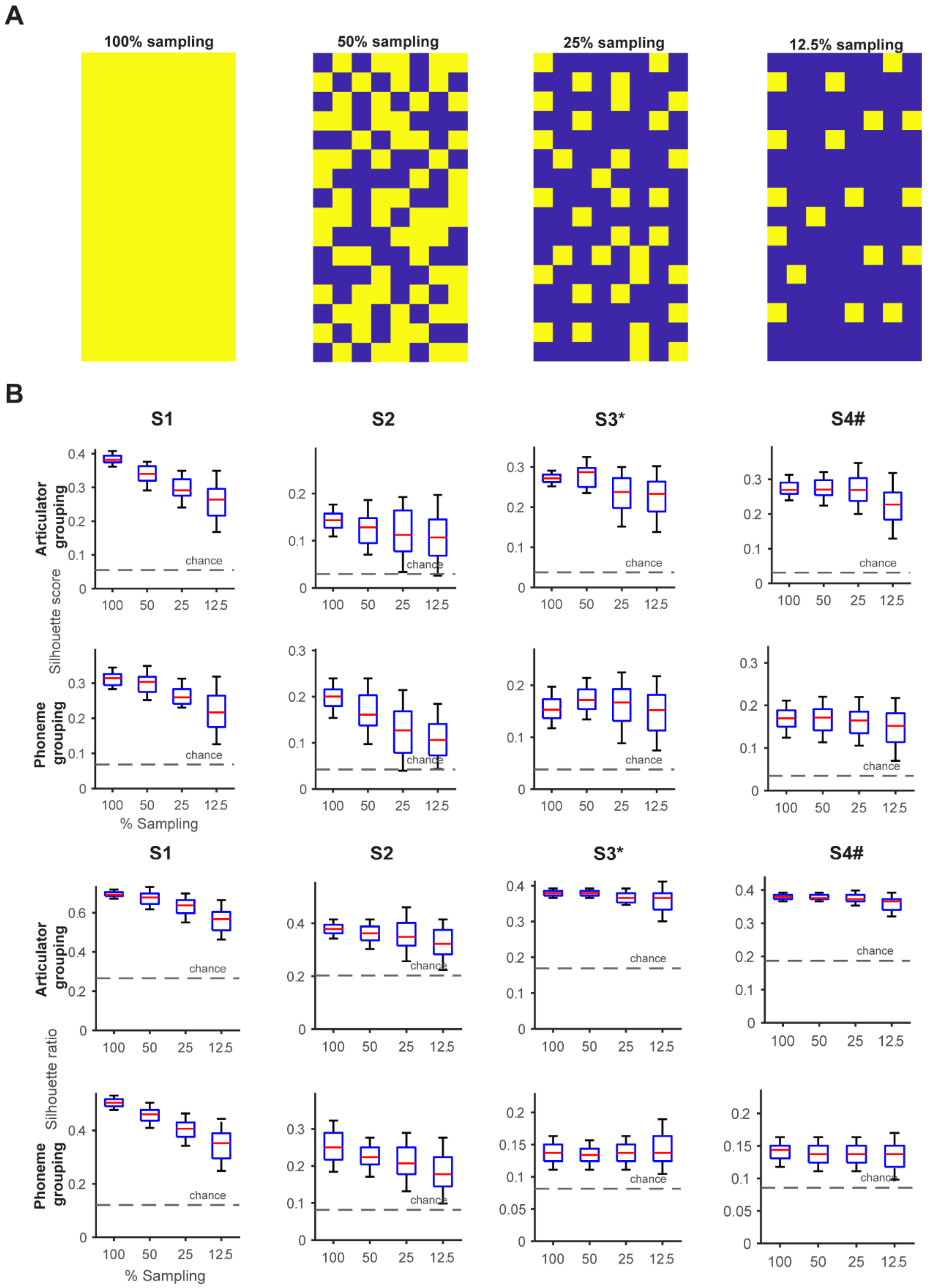


**Supplementary Figure 4**. **High resolution recording enables robust cortical state-space for phoneme articulators in SMC.** **A)** Demonstration of electrode subsampling using Poisson-disc method. Subsampled electrodes are uniformly distributed across the array. This method of uniform subsampling provides valid low-resolution simulation when compared to random sampling of electrodes. **B)** Silhouette score and silhouette ratio measures clustering of utterance trials in the cortical state-space (SVD – tSNE) with respect to either articulatory or phoneme grouping (* - low SNR recording, # - completed 1 block only). Spatial subsampling of electrodes using Poisson-disc sampling at fixed spatial coverage deforms the cortical-state-space for all subjects, as measured by a decrease in silhouette values. Each distribution contains state-space evaluations from 50 electrode sub-samplings. The red lines and blue boxes indicate the median and 25/75$^{th}$ percentile, and the dotted lines indicate the full range of the distributions. High-resolution cortical sampling enables accurate capturing of speech articulatory features in SMC.

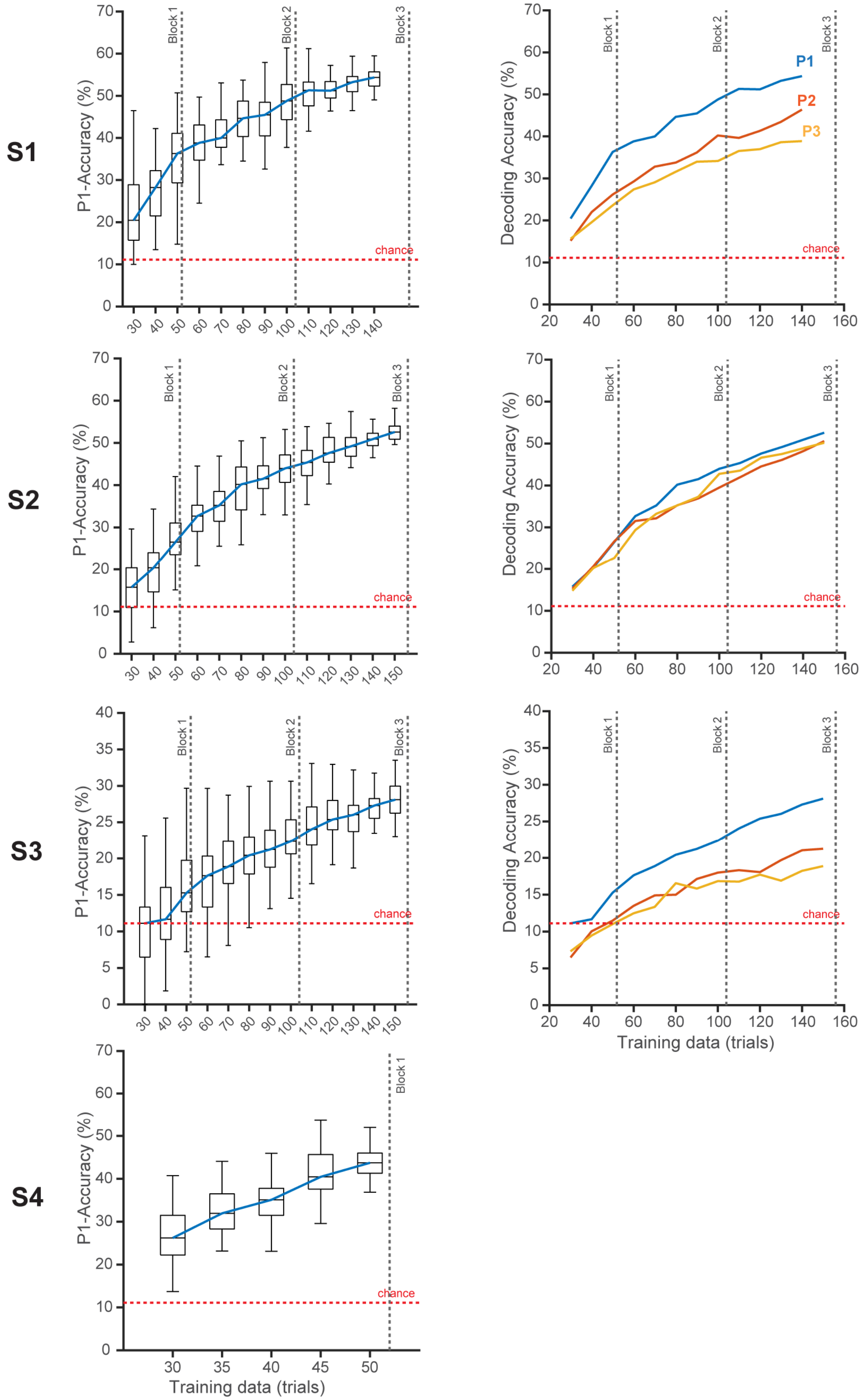


**Supplementary Figure 5**. **Effect of training duration on decoding performance for all four subjects**. To investigate the impact of training duration, decoding performance is calculated on subsampled trials (50 samples) by maintaining equal distribution of phoneme trials in each sampling. The decoding performance (**left column**) increases with increased data for all subjects, as indicated my median decoding accuracy. The vertical dotted lines indicate the number of trials the patients would have completed by the end of an experimental block. These performance trends were stable across all phoneme positions as indicated my median decoding performance for P1, P2, and P3 (**right column**). Subject S4 completed one-block of speech experiment, thereby, limiting the number of trials required to perform subsampling analysis for P2 and P3.

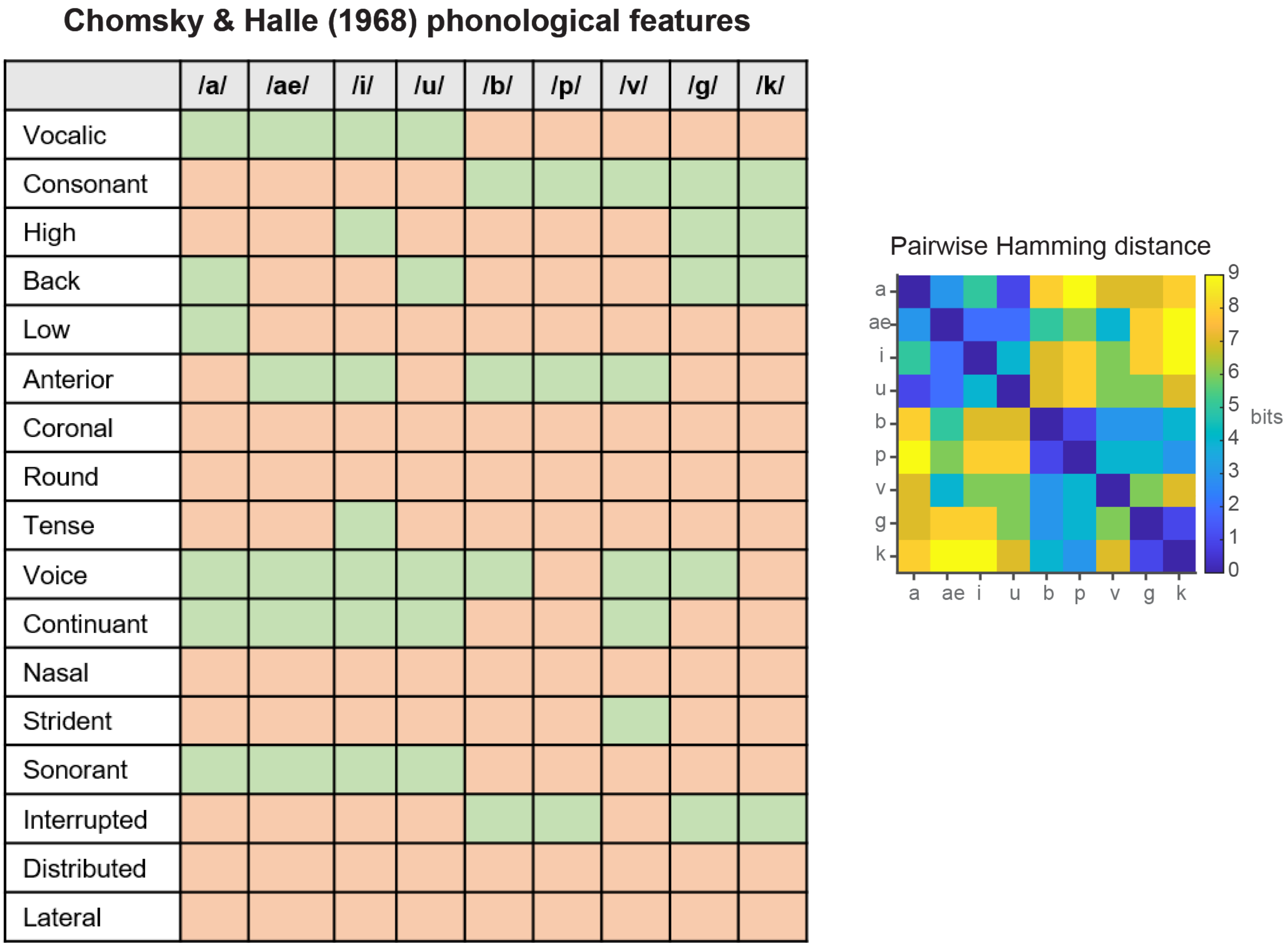


**Supplementary Figure 6**. **Chomsky & Halle (1968) phonological features for phonemes used in our study**. Each phoneme is identified by 17-bit binary feature vector (green = 1, red = 0) to characterize phonological articulatory features. A pairwise distance metric was computed between all possible phoneme pairs using Hamming distance. The resultant phonological distance values were lower for phonemes with similar phonological features and vice-versa (e.g. the phoneme /p/ differed from /b/ by 1 bit, and from /g/ by 5 bits).

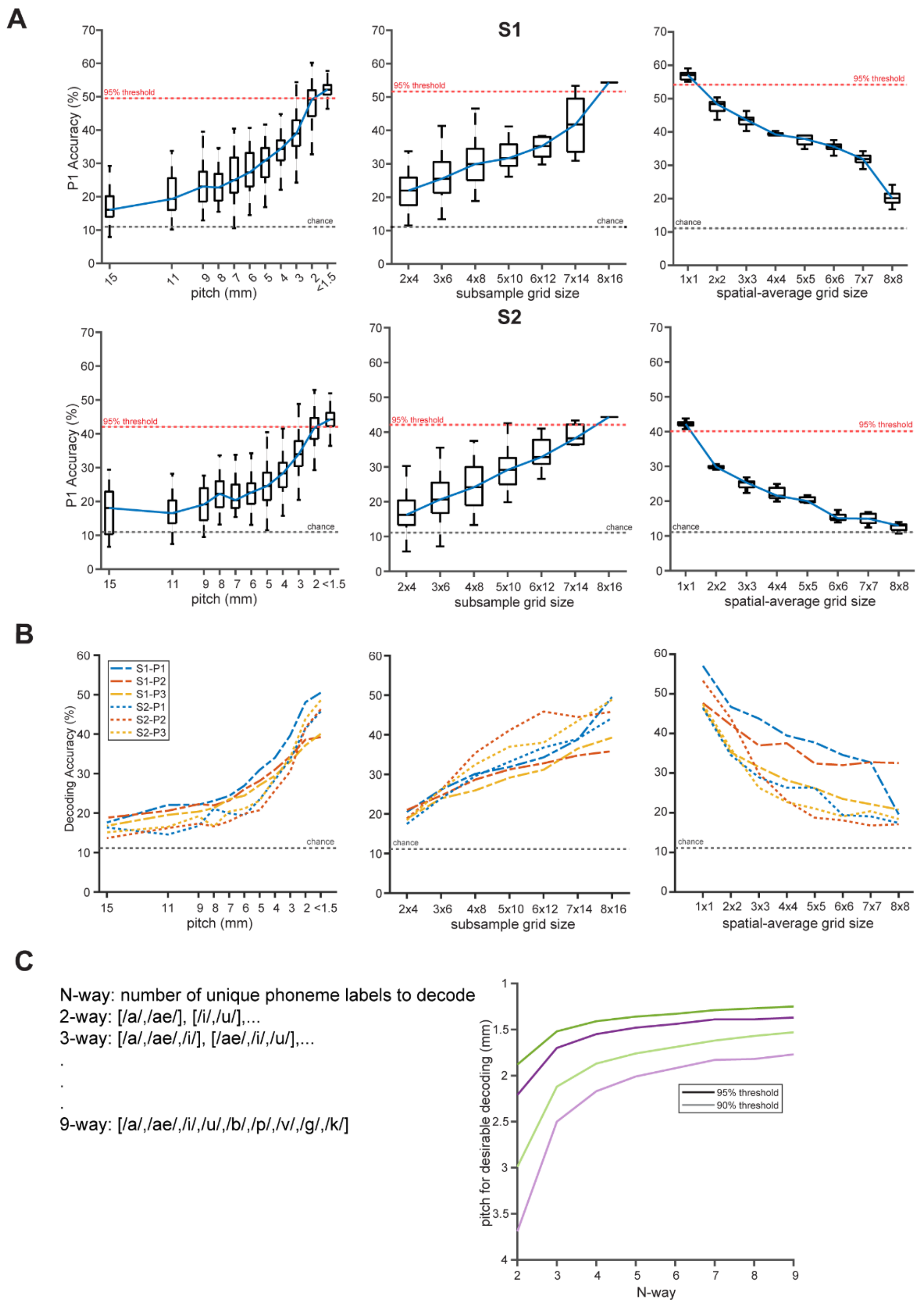


**Supplementary Figure 7 Effect of spatial-resolution, coverage, and micro-scale contact size on decoding performance for both S1 and S2. A) Left column:** To investigate spatial resolution, P1- decoding accuracy was calculated for subsampled electrodes (50 samples using Poisson disc sampling) at maintained coverage. Subsampled electrode populations with resolutions less than 1.5 mm crossed the 95% threshold. The decoding performance is improved by 17% and 29% with respect to 4-mm and 10-mm pitch, respectively. **Middle column:** To test for spatial coverage, decoding performance was assessed each rectangular subgrid at each possible array location. P1 accuracy increased with increasing coverage for both S1 and S2, and the maximum performance was achieved at the widest coverage possible. **Right column:** To examine the effect of micro-scale sampling, P1 accuracy was calculated on spatially averaged neural data with smoothing windows of varying grid-sizes. Decoding performance was maximal with the micro-scale contact size (200 μm) and decreased with increased spatial smoothing (i.e. increased synthetic contact size). The 95% max-accuracy thresholds and theoretical chance boundaries were drawn separately for each subject. **B)** Median decoding scores indicate that the influence of spatial-resolution, coverage, and micro-scale contact is stable for phonemes across all positions for both S1 and S2. **C)** Effect of spatial-resolution on the ability to decode the number of unique phoneme labels (N) for both S1 (violet) and S2 (green). Spatial resolution required to achieve 90-95% of maximum decoding performance, increased with increase in N. Thus, the analysis demonstrated the necessity of spatial-resolution, spatial coverage, and micro-scale sampling for accurate speech decoding.

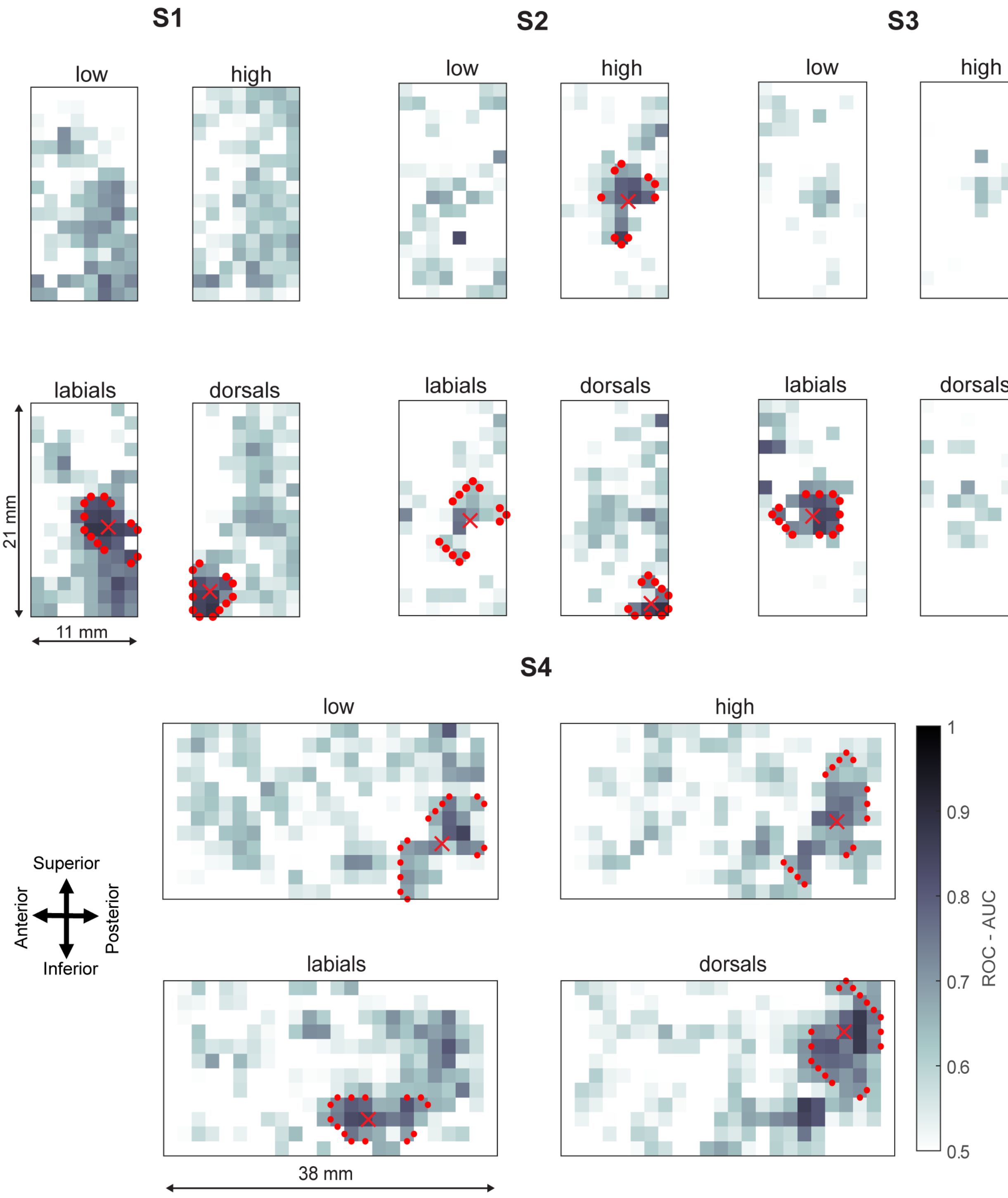


**Supplementary Figure 8**. **Articulator encoding maps for high-resolution neural recordings**. Univariate decoding of four articulatory features (low vowels, high vowels, labial consonants, dorsal consonants) identified articulatory tuning for each electrode. The spatial arrangement of resultant AUC values shows articulatory feature maps for all subjects. These maps showed distinct spatial clusters within the array, as identified by red contours, which identify the 90$^{th}$ percentile of AUC values across all articulators (colored marker x indicates the corresponding centroids).

**Supplementary Table 1:**

**Clinical Summary of patients: Intra-operative awake surgery**

| Patient | Age | Sex | Diagnosis | Electrode type/ No. of recording electrodes (coverage) | Electrode material | Number of electrodes with significant HG activation in SMC (estimated coverage) |
|---|---|---|---|---|---|---|
| S1 | 61 | M | Parkinson's | μECoG: 128 (231 $mm^2$) | Gold | 111 (199 $mm^2$) |
| S2 | 63 | M | Parkinson's | μECoG: 128 (231 $mm^2$) | Platinum-Iridium | 111 (199 $mm^2$) |
| S3 | 62 | M | Parkinson's | μECoG: 128 (231 $mm^2$) | Platinum-Iridium | 60 (108 $mm^2$) |
| S4 | 26 | F | Tumor resection | μECoG: 256 (798 $mm^2$) | Platinum-Iridium | 149 (440 $mm^2$) |

**Clinical Summary of patients: Pre-operative epileptic monitoring**

| Patient | Age | Sex | Diagnosis | Electrode type/ Total No. of recording electrodes | Electrode spacing s (mm) | Number of electrodes with significant HG activation in SMC (estimated coverage) |
|---|---|---|---|---|---|---|
| D6 | 28 | M | Epilepsy | Ad-Tech 64 channel ECoG: 96 | 10 | 12 (*1200 $mm^2$) |
| D7 | 34 | M | Epilepsy | Ad-Tech 64 channel ECoG: 102 | 10 | 10 (*1000 $mm^2$) |
| D24 | 38 | F | Epilepsy | Ad-Tech 48 channel ECoG: 52 | 10 | 3 (*300 $mm^2$) |
| D26 | 23 | F | Epilepsy | Ad-Tech 48 channel ECoG: 60 | 10 | 1 (*100 $mm^2$) |
| D35 | 21 | M | Epilepsy | PMT SEEG: 174 | 3.5 | 4 (**38.4 $mm^2$) |
| D40 | 27 | M | Epilepsy | PMT SEEG: 188 | 3.5 | 2 (**19.2 $mm^2$) |
| D52 | 24 | F | Epilepsy | PMT SEEG: 194 | 3.5 | 5 (**48.1 $mm^2$) |
| D55 | 30 | F | Epilepsy | PMT SEEG: 190 | 3.5 | 6 (**57.7 $mm^2$) |
| D59 | 8 | F | Epilepsy | PMT SEEG: 184 | 3.5 | 4 (**38.5 $mm^2$) |
| D60 | 40 | F | Epilepsy | PMT SEEG: 252 | 3.5 | 20 (**192.4 $mm^2$) |
| D70 | 41 | F | Epilepsy | Ad-Tech SEEG: 202 | 5 | 4 (**78.5 $mm^2$) |

* Estimated coverage for ECoG: $A = n * s^2$;

** Estimated coverage for SEEG: $A = n * \pi * \left(\frac{s}{2}\right)^2$; n – number of electrodes, s – electrode spacing

**Supplementary Table 2: Stimulus labels for speech repetition task**

| CVC | VCV |
|---|---|
| /bab/ | /abae/ |
| /baek/ | /abi/ |
| /bak/ | /aeba/ |
| /bup/ | /aebi/ |
| /gab/ | /aebu/ |
| /gaeb/ | /aega/ |
| /gaev/ | /aeka/ |
| /gak/ | /aepi/ |
| /gav/ | /aka/ |
| /gig/ | /aku/ |
| /gip/ | /ava/ |
| /gub/ | /avae/ |
| /kab/ | /ibu/ |
| /kaeg/ | /ika/ |
| /kub/ | /ikae/ |
| /kug/ | /ipu/ |
| /paek/ | /iva/ |
| /paep/ | /ivu/ |
| /paev/ | /uba/ |
| /puk/ | /uga/ |
| /pup/ | /ugae/ |
| /vaek/ | /ukae/ |
| /vaeg/ | /upi/ |
| /vip/ | /upu/ |
| /vug/ | /uvae/ |
| /vuk/ | /uvi/ |

**Supplementary Table 3: Neural data duration for each intraoperative patient**

| Patient | Blocks/Trials | Total spoken duration (minutes) | Number of correct trial repetitions (n) | Neural data duration (minutes; 1s per trial) T = n * 1/60 | Neural data duration to train SVD-LDA (minutes; 20-fold CV) 0.95 * T | Neural data duration to train seq2seq RNN (minutes; 80% training held out) 0.8 * T |
|---|---|---|---|---|---|---|
| S1 | 3/156 | 1.04 | 149 | 2.48 | 2.36 | 1.98 |
| S2 | 3/156 | 1.15 | 152 | 2.53 | 2.4 | 2.02 |
| S3 | 3/156 | 1.45 | 153 | 2.55 | 2.42 | N.A. |
| S4 | 1/52 | 0.47 | 52 | 0.87 | 0.83 | N.A. |

N.A. – not applicable

**Supplementary Table 4: Articulator coverage and tuning**

| Subjects | | Low | High | Labials | Dorsal |
|---|---|---|---|---|---|
| S1 (16 x 8) | coverage ($mm^2$) | ~ | ~ | 72.5 | 17.7 |
| | mean AUC | ~ | ~ | 0.9 | 0.9 |
| | Centroid coordinates (X, Y) | ~ | ~ | (9.8, 6.2) | (14.7, 1.7) |
| S2 (16 x 8) | | ~ | 26.5 | 28.3 | 10.6 |
| | | ~ | 0.88 | 0.82 | 0.91 |
| | | ~ | (9.1, 5.4) | (9.2, 5.6) | (15.4, 7.2) |
| S3 (16 x 8) | | ~ | ~ | 47.8 | ~ |
| | | ~ | ~ | 0.81 | ~ |
| | | ~ | ~ | (8.9, 4.6) | |
| S4 (12 x 24) | | 79.88 | 85.8 | 56.2 | 79.88 |
| | | 0.85 | 0.8 | 0.84 | 0.88 |
| | | (8.5, 20.3) | (7.2, 20.3) | (9.8, 15.6) | (4.6, 21.1) |

~ spatial clustering not observed

**Supplementary Table 5: Hyperparameters for seq2seq based encoder-decoder model**

| Hyperparameters | S1 | S2 |
|---|---|---|
| Number of convolutional filters | 100 | 90 |
| RNN units | 800 | 900 |
| *l2* penalty | 1e-6 | 1e-5 |
| 1D convolutional filter length | 10 | |
| Number of training epochs | 800 | |
| Learning rate | 1e-3 | |
| Optimization | Adam | |

**Supplementary Table 6: Example decoded phoneme sequences using seq2seq based encoder-decoder model**

## S1

| True | Predicted |
|---|---|
| '/vuk/' | '/vuk/' |
| '/aeka/' | '/aeka/' |
| '/gip/' | '/gip/' |
| '/gub/' | '/gub/' |
| '/puk/' | '/pup/' |
| '/baek/' | '/vaek/' |
| '/baek/' | '/aeaek/' |
| '/abae/' | '/bbae/' |
| '/vuk/' | '/puk/' |
| '/uba/' | '/aeba/' |
| '/uvae/' | '/ubae/' |
| '/bup/' | '/vup/' |
| '/vug/' | '/vuk/' |
| '/vaek/' | '/vak/' |
| '/bak/' | '/aak/' |
| '/ika/' | '/aeaa/' |
| '/uvi/' | '/uak/' |
| '/gaeb/' | '/kab/' |
| '/uga/' | '/aeba/' |
| '/gaev/' | '/kaeg/' |
| '/uba/' | '/upu/' |
| '/ava/' | '/aeba/' |
| '/paev/' | '/vaek/' |
| '/aka/' | '/aega/' |
| '/upi/' | '/vui/' |
| '/aku/' | '/aga/' |
| '/paep/' | '/bab/' |
| '/gaeb/' | '/ipi/' |
| '/upu/' | '/aba/' |
| '/paek/' | '/aka/' |

## S2

| True | Predicted |
|---|---|
| '/gaev/' | '/gaev/' |
| '/gip/' | '/gip/' |
| '/gak/' | '/gak/' |
| '/ibu/' | '/ibu/' |
| '/abae/' | '/abae/' |
| '/kab/' | '/kab/' |
| '/kub/' | '/kub/' |
| '/bab/' | '/baa/' |
| '/vug/' | '/gug/' |
| '/baek/' | '/baep/' |
| '/gig/' | '/gib/' |
| '/upi/' | '/ipi/' |
| '/vip/' | '/pip/' |
| '/gip/' | '/pip/' |
| '/ava/' | '/aba/' |
| '/aka/' | '/aeka/' |
| '/paek/' | '/vaek/' |
| '/gab/' | '/bab/' |
| '/uvae/' | '/aevae/' |
| '/upu/' | '/uku/' |
| '/uga/' | '/aeka/' |
| '/avae/' | '/aebae/' |
| '/ipu/' | '/ubu/' |
| '/gav/' | '/gub/' |
| '/paev/' | '/aeaeae/' |
| '/puk/' | '/vug/' |
| '/aeka/' | '/aebae/' |
| '/aku/' | '/ubu/' |
| '/uvae/' | '/aba/' |
| '/ipu/' | '/ugae/' |
| '/iva/' | '/aeaeae/' |
